# Diversity of trypanosomatids in small mammals from a forest fragment at the wildlife-urban interface in southeastern Brazil: implications for public health surveillance

**DOI:** 10.1101/2025.08.08.669276

**Authors:** Amanda Caroline Corrêa Madureira, Ana Paula Isnard, Ana Cristina Vianna Mariano da Rocha Lima, Débora Cristina Capucci, Anna Luiza Moreira Martins, Letícia Gracielle Tôrres de Miranda Estevam, Mariana Lourenço Freire, Soraia de Oliveira Silva, Leonardo Barbosa Koerich, Grasiele Caldas D’Avila Pessoa, Nelder de Figueiredo Gontijo, Daniel Moreira de Avelar, Rita de Cássia Moreira de Souza, Lileia Gonçalves Diotaiuti, José Dilermando Andrade Filho, Felipe Dutra-Rêgo

## Abstract

This study investigated the diversity of trypanosomatids infecting small mammals in a fragmented forest landscape in southeastern Brazil, aiming to clarify their role in transmission cycles and assess potential public health risks. Eighteen individuals from three species - *Didelphis albiventris* (n = 8), *Marmosops incanus* (n = 4), and *Nectomys squamipes* (n = 6) - were captured in the Mata da Tapera Municipal Natural Park (MT), located within the Espinhaço Range Biosphere Reserve, Minas Gerais State. Xenodiagnosis using *Lutzomyia longipalpis* and *Rhodnius neglectus*, followed by 18S rDNA nested PCR, enabled the detection of *Leishmania infantum*, *L. braziliensis*, and *Trypanosoma cruzi* DTU TcI in insects fed on naturally infected hosts. Tissue and organ samples were also analyzed by in vitro culture and molecular diagnostics. *Leishmania infantum* was identified in *D. albiventris* (n = 2), *L. braziliensis* in *M. incanus* (n = 1), *T. cruzi* TcI in *D. albiventris* (n = 2), and *Trypanosoma lainsoni* in both *D. albiventris* and *N. squamipes* (n = 1 each), the latter representing the first record of this parasite in that rodent species. This integrated approach provides a valuable model for zoonotic surveillance at the wildlife-urban interface, enhancing diagnostic sensitivity and species identification. These findings confirm the presence of zoonotic trypanosomatids in a highly anthropized landscape and highlight transmission risks to humans and domestic animals, particularly in an area with active ecotourism, underscoring the need for targeted surveillance within a One Health framework.

**Author summary:** In many parts of Brazil, forested areas are getting smaller and more fragmented due to urban growth. These changes increase the contact between wild animals, domestic animals, and humans, creating new opportunities for diseases to spread. In this study, we examined small wild mammals living in a forest fragment close to human settlements in southeastern Brazil to find out if they carried parasites that can also infect humans. We used a combination of techniques, including allowing laboratory-raised insects to feed on these animals, to detect the presence of different parasites. We found that these mammals were naturally infected with species of *Leishmania* and *Trypanosoma*, including those that cause leishmaniasis and Chagas disease in humans. We also detected *Trypanosoma lainsoni*, a lesser-known parasite, for the first time in one rodent species. Our findings suggest that small mammals living in areas where forests meet cities may play an important role in maintaining and spreading parasites that affect public health. This study shows the importance of monitoring wildlife in changing environments and supports the idea that human, animal, and environmental health are deeply connected.

## Introduction

Diverse environmental factors influence the transmission, incidence, and dynamics of vector-borne diseases by directly or indirectly affecting pathogen spread or by altering the life cycles of vectors and reservoirs, ultimately shaping their distribution and abundance [1,2]. Forest disturbance and fragmentation can lead to habitat loss and changes in blood meal availability for vectors, emerging as critical drivers in the transmission dynamics of leishmaniasis and Chagas disease [3,4]. Transitional zones between wild and urban environments are often subject to biodiversity loss, favoring generalist species, such as rodents and marsupials, that can act as pathogen reservoirs [5–7]. These ecological shifts intensify interactions among humans, domestic animals, and wildlife, creating favorable conditions for spillover and spillback events [8].

In southeastern Brazil, where rapid urban expansion overlaps remnants of the Atlantic Forest and *Cerrado* biomes [9,10], peri-urban forest fragments represent key ecological interfaces between human settlements and wildlife. In these transitional landscapes, synanthropic mammals, particularly rodents and marsupials, serve as both reservoirs and sentinels for a broad diversity of trypanosomatids, from non-pathogenic to medically important species. These hosts may contribute to parasite amplification across heterogeneous environments and sustain transmission cycles that are often overlooked due to the cryptic nature of host-parasite associations [11–13].

Despite their potential role in maintaining parasite circulation, the structure of host-parasite interactions and their ability to sustain enzootic or zoonotic cycles in disturbed peri-urban forests remain insufficiently understood. While classical transmission areas have been extensively studied, fewer investigations have focused on transitional landscapes subjected to intense anthropogenic pressure [6,14].

Here, we investigated the diversity of trypanosomatids infecting small mammals in a forest fragment located at the wildlife-urban interface in southeastern Brazil. This region has previously reported autochthonous cases of canine visceral leishmaniasis (unpublished data, State Health Department of Minas Gerais), and phlebotomine sand flies carrying *Leishmania* DNA have also been recorded [15]. In addition, *Panstrongylus megistus*, one of the primary vectors of Chagas disease in Brazil, is also present in the area [16,17]. By characterizing infections in small mammals and assessing their potential to infect known vector species, we provide insights into the ecological dynamics of trypanosomatid transmission in transitional environments. Our findings highlight the role of small mammal communities as reservoirs within complex transmission networks and underscore the epidemiological implications of parasite circulation in areas under increasing anthropogenic pressure.

## Material and Methods

### Ethical statements

This study was approved by the Animal Ethics Committee (CEUA) of the Oswaldo Cruz Foundation (protocol number LW-3/23) and authorized by the Authorization and Information on Biodiversity System (SISBIO; permit number 86644-1). The collection of biological material from mammals and access to genetic resources were conducted in compliance with Brazilian regulations. The study was registered in the National System for the Management of Genetic Heritage and Associated Traditional Knowledge (SisGen) under registration number A47F256.

### Study area and sample collection

The Mata da Tapera Municipal Natural Park (MT), located in the Serra do Cipó district of Minas Gerais State (19°19’58.7’S 43°36’52.5’W), is part of a network of protected areas known as the “*Cipó Mosaic*”, a UNESCO-recognized site and part of the Espinhaço Range Biosphere Reserve. Although it is an anthropized forest fragment with close interactions among humans, domestic animals, and wildlife, the park holds significant socio-environmental value due to its natural springs and remnants of the Brazilian *Cerrado*.

Between March 2023 and September 2024, six small mammal sampling campaigns were conducted in the MT (Fig 1), encompassing both the rainy and dry seasons (three campaigns per season). Small mammals were captured using galvanized steel traps, Tomahawk model, baited with a mixture of fruit and cod liver oil emulsion or bacon. Specimens were identified following the taxonomic keys proposed for Brazilian mammals [18,19], and all specimens were deposited in the Mammal Collection of the Natural Sciences Museum, Pontifical Catholic University of Minas Gerais (PUC-Minas).

**Fig 1:**
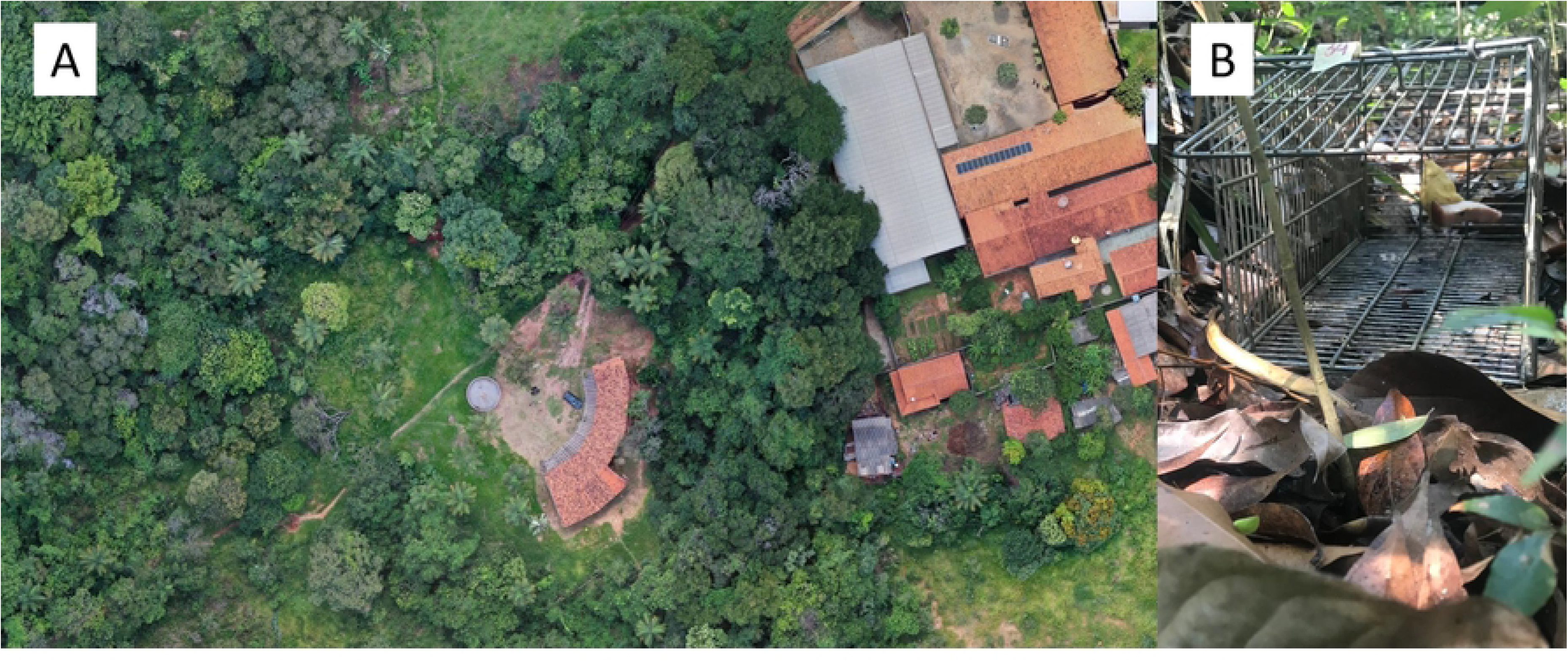
Small mammal collection site within Mata da Tapera Municipal Natural Park (MT), located in the Serra do Cipó district, Minas Gerais State, Brazil. (A) General view of the MT area, showing its proximity to peridomestic environments. (B) Steel traps (Tomahawk model) baited with a mixture of fruit and cod liver oil emulsion or bacon, used for small mammal captures. Images licensed under CC BY, with permission from Felipe Dutra Rêgo, copyright 2025.

Following species identification, animals were anesthetized with a combination of xylazine (10 mg/kg) and ketamine (200 mg/kg) for xenodiagnosis and intracardiac blood collection. Subsequently, euthanasia was performed by anesthetic overdose to enable the collection of organs and tissues fragments for the investigation of natural trypanosomatid infections through both *in vitro* culture and molecular detection using nested PCR targeting the 18S rDNA.

### Determination of infectivity via xenodiagnosis

Anesthetized small mammals were positioned in dorsal recumbency for xenodiagnosis procedures using *Lutzomyia longipalpis* (Piauí, Brazil), obtained from the phlebotomine sand fly colony maintained at the Laboratory of Physiology of Hematophagous Insects, Federal University of Minas Gerais (UFMG), and *Rhodnius neglectus* (Tocantinópolis, Tocantins, Brazil), from the triatomine colony of the René Rachou Institute, Oswaldo Cruz Foundation (Fiocruz Minas).

Insects were placed in ventilated plastic containers covered with permeable mesh and positioned over the abdominal region and limbs of the animals to allow blood feeding for approximately 30 minutes (Fig 2). For each mammal, approximately 100 female and 20 male *L. longipalpis*, and 10 third- or fourth-instar *R. neglectus* nymphs were used. After feeding, engorged sand flies were transferred to insulated containers with continuous monitoring of temperature and humidity (25 °C, 80% RH), and subsequently maintained in the insectary of the Medical Entomology Group at IRR/Fiocruz Minas. Sand flies were provided with a 15% glucose solution *ad libitum* during the holding period. Triatomines were maintained in an incubator under controlled conditions (27 °C, 60% RH, 12:12 h light/dark cycle) and were fed on chicken blood 10 days after xenodiagnosis.

**Fig 2:**
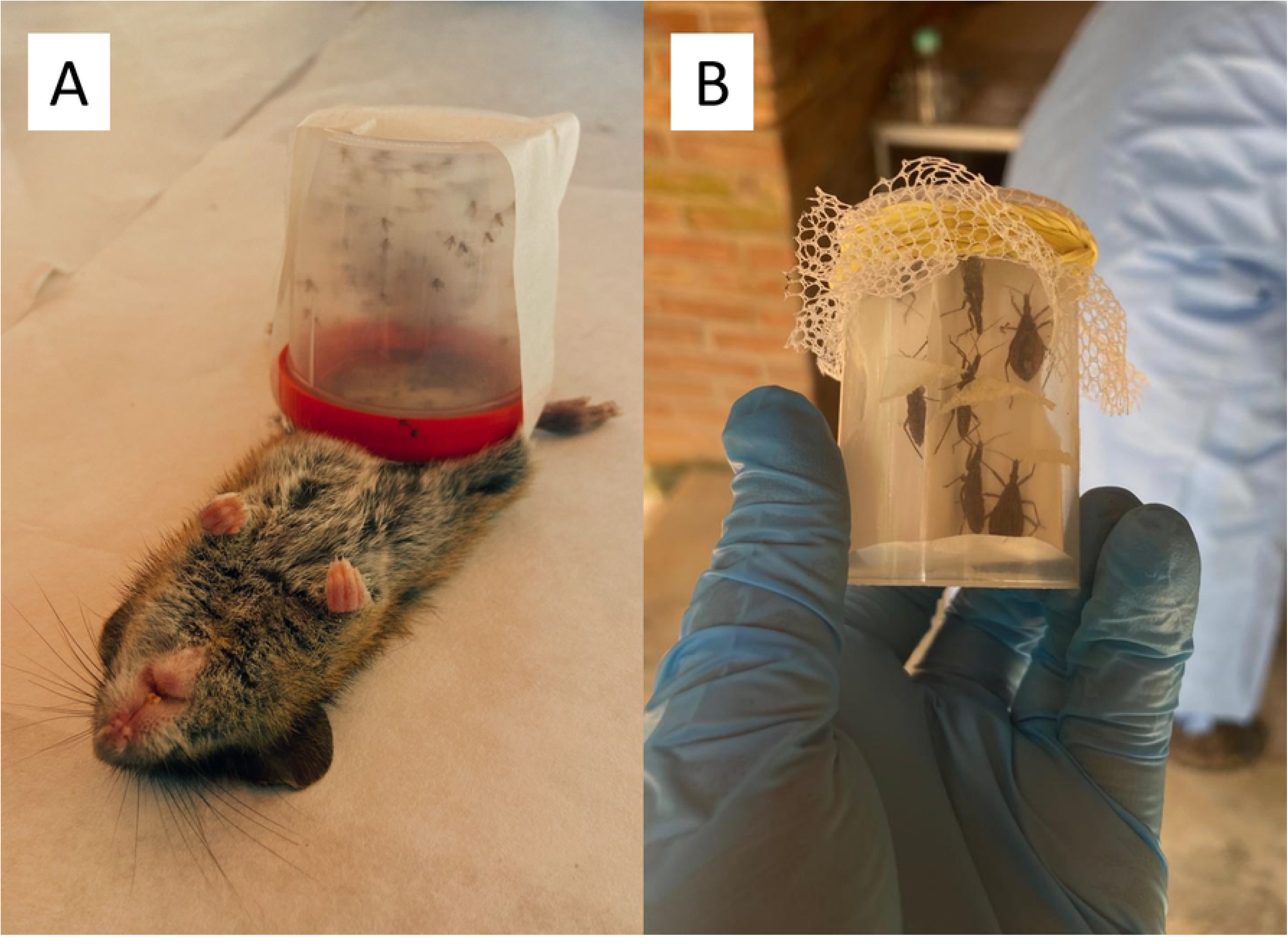
Field xenodiagnosis of wild and synanthropic small mammals using laboratory-reared *Lutzomyia longipalpis* and *Rhodnius neglectus* in Mata da Tapera Municipal Natural Park (MT), Serra do Cipó district, Minas Gerais State, Brazil. (A) *Nectomys squamipes* under anesthesia during xenodiagnosis with *Lutzomyia longipalpis*. **(B)** Fourth-instar nymphs of *R. neglectus* used in the procedure. Animals were maintained under sedation with xylazine (10 mg/kg) and ketamine (200 mg/kg) for approximately 30 minutes during xenodiagnostic exposure.

*Lutzomyia longipalpis* specimens were euthanized five days post-feeding and processed for molecular detection of *Leishmania* via 18S rDNA nested PCR. *R. neglectus* individuals were assessed 30 days post-exposure. For parasitological examination, feces were obtained by gentle abdominal compression and placed on a microscope slide with a drop of 1× PBS. A coverslip was added, and the sample was examined under a binocular light microscope at 400× magnification. All visual fields were systematically screened for the presence of *T. cruzi*.

### Natural infection of small mammals by Trypanosomatids

Whole blood aliquots were immediately inoculated in the field into biphasic NNN/LIT culture medium. Tissue fragments (ear skin, tail skin, heart, spleen, and liver) were stored at 4 °C for at least 72 hours in phosphate-buffered saline (PBS 1×, pH 7.2), supplemented with antibiotics (penicillin, 5,000 U/mL; streptomycin, 5,000 µg/mL) and an antifungal agent (5-fluorocytosine, 50 mg/L). After homogenization under a laminar flow hood, tissues were inoculated into biphasic NNN/LIT medium supplemented with 20% sterile, heat-inactivated fetal bovine serum and antibiotics, as previously described.

Cultures were maintained in individually labeled 15 mL conical tubes and incubated at 25 °C (±1 °C). Each culture was examined weekly under light microscopy for the presence of flagellated forms and subcultured into fresh medium as needed, for up to three months. Cultures exhibiting fungal or bacterial contamination were discarded based on macroscopic and microscopic evaluation. Positive cultures were centrifuged at 2,500 × g for 10 minutes, and the resulting parasite pellet was subjected to DNA extraction using the Gentra Puregene Tissue Kit (Qiagen), followed by molecular identification using V7V8-targeted nested PCR [20,21].

### Molecular detection of trypanosomatids

Fragments of ear skin, tail skin, heart, spleen, liver, lung, intestine, and lymph nodes were collected for molecular diagnostics. All samples were individually labeled, preserved in RNAlater (Invitrogen), and stored at −20 °C until DNA extraction. Total DNA was extracted using the Gentra Puregene Tissue Kit (Qiagen), following the manufacturer’s instructions, and eluted in 50 µL of elution buffer.

To assess DNA integrity and detect potential PCR inhibitors, whole blood samples were subjected to amplification of the mitochondrial cytochrome *b* gene using primers cyt b1 (5′-CCATCCAACATCTCAGCATGATGAAA-3′) and cyt b2 (5′-GCCCCTCAGAATGATATTTGTCCTCA-3′) [22]. For tissue and organ samples, DNA quality was evaluated by amplifying a 70 bp fragment of the γ-actin gene using primers γactin-For (5′-ACAGAGAGAAGATGACGCAGATAATG-3′) and γactin-Rev (5′-GCCTGAATGGCCACGTACA-3′) [23].

Trypanosomatid detection was performed using a nested PCR targeting the V7V8 variable region of the 18S rDNA [20,21]. Each PCR assay included a negative control (no DNA template) and positive controls containing 10 ng of DNA from *Leishmania amazonensis* (IFLA/BR/67/PH8) and *Trypanosoma cruzi* (DTU TcI).

PCR-positive amplicons were purified using either the QIAquick PCR Purification Kit or the QIAquick Gel Extraction Kit (Qiagen), depending on product resolution. Sequences were analyzed using FinchTV (Geospiza, Inc.), aligned in MEGA X [24], and compared with reference sequences from GenBank using BLAST. Sequences generated in this study were deposited in GenBank under accession numbers PV688719-PV688727.

## Results

A total of 18 small mammals were captured, representing three species: *Nectomys squamipes* (n = 9; 50%), *Didelphis albiventris* (n = 5; 27.8%), and *Marmosops incanus* (n = 4; 22.2%). Most individuals (72%) were captured in June 2023 (n = 5; 28%), followed by August 2023 and September 2024 (n = 4; 22% each). One *D. albiventris* female, which was carrying dependent offspring, was released at the capture site and excluded from the analyses (Table 1). No cutaneous lesions were observed during clinical examination of any individual in the field.

**Table 1:** Small mammal species captured in Mata da Tapera Municipal Natural Park (Minas Gerais, Brazil), by collection month.

| Species | Collection month |  |  |  |  |  | Total (%) |
| --- | --- | --- | --- | --- | --- | --- | --- |
|  | Mar/23 | Jun/23 | Aug/23 | Sep/23 | Jan/24 | Sep/24 |  |
| *Didelphis albiventris* | 2 | - | 1* | - | 1 | 1 | **5 (28)** |
| *Nectomys squamipes* | - | 3 | 1 | 2 | - | 3 | **9 (50)** |
| *Marmosops incanus* | - | 2 | 02 | - | - | - | **4 (22)** |
| **Total (%)** | **2 (11)** | **5 (28)** | **4 (22)** | **2 (11)** | **1 (6)** | **4 (22)** | **18 (100)** |

\*One specimen of *Didelphis albiventris* was released due to the presence of offspring.

Two *M. incanus* specimens died before xenodiagnosis and were not included in the assay. A total of 397 *Lutzomyia longipalpis* females that fed on the remaining 15 animals were individually analyzed by nested PCR. The number of females screened per host ranged from 10 to 46, depending on survival five days post-feeding. Four females tested positive for *Leishmania*: *L. infantum* was detected in sand flies fed on *N. squamipes* (n = 2) and *D. albiventris* (n = 1), while *L. braziliensis* was identified in one fly fed on *M. incanus* (Fig 3).

**Fig 3:**
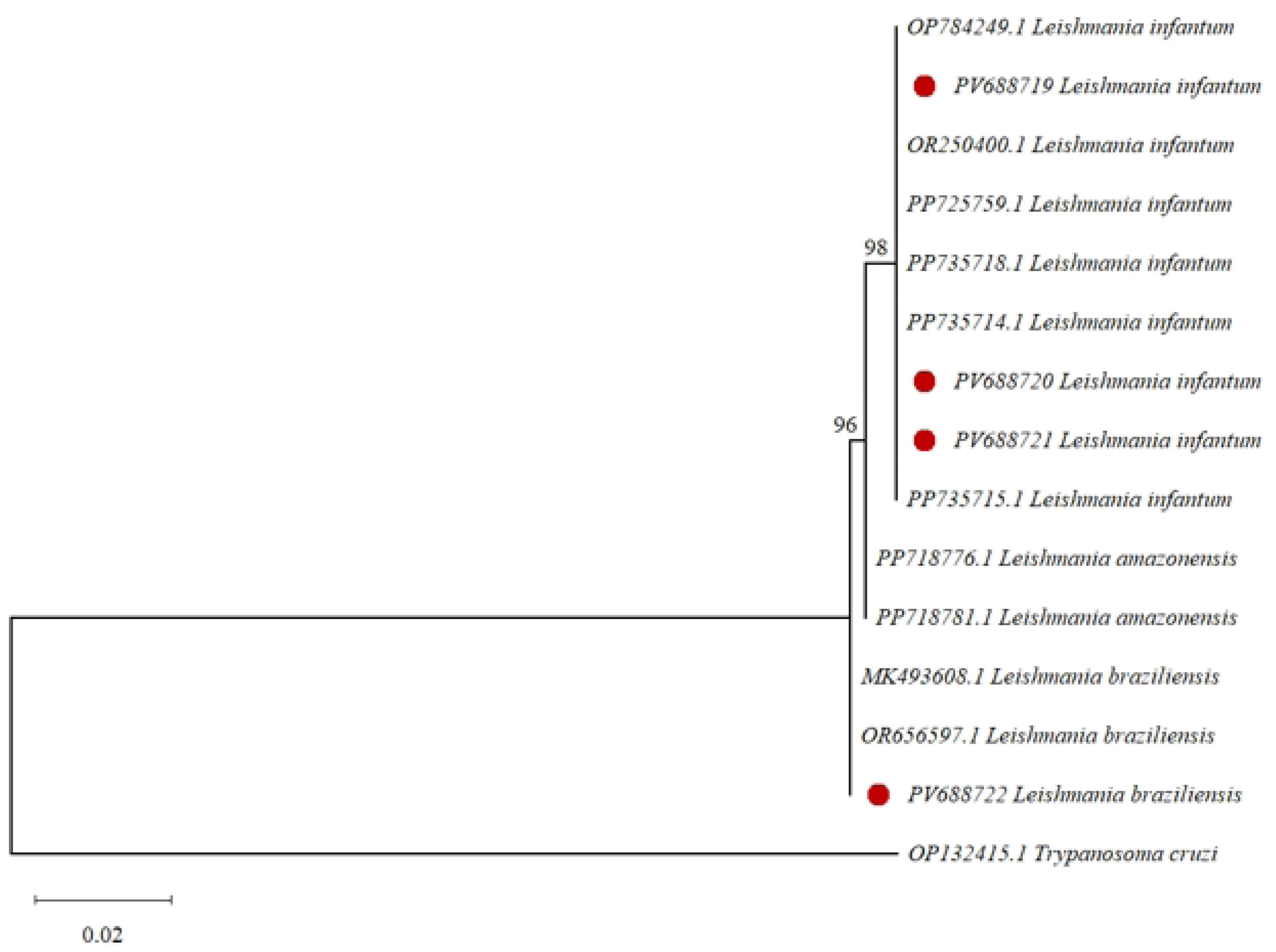
Phylogenetic tree of *Leishmania* spp. based on isolates from small mammals captured in Mata da Tapera Municipal Natural Park (MT), Minas Gerais, Brazil. The tree was inferred from 18S SSU rDNA sequences using the Tamura-Nei model with gamma-distributed rate variation among sites (TN+G). Bootstrap values (1,000 replicates) are shown at the nodes. Sequences obtained in this study are marked with red circles. The scale bar indicates the number of nucleotide substitutions per site.

Regarding xenodiagnosis using triatomines, *Trypanosoma cruzi* DTU TcI was detected in two *D. albiventris* individuals. All nymphs exposed to animal ID02 tested positive, whereas only two out of five nymphs were positive for ID16 (Fig 4).

**Fig 4:**
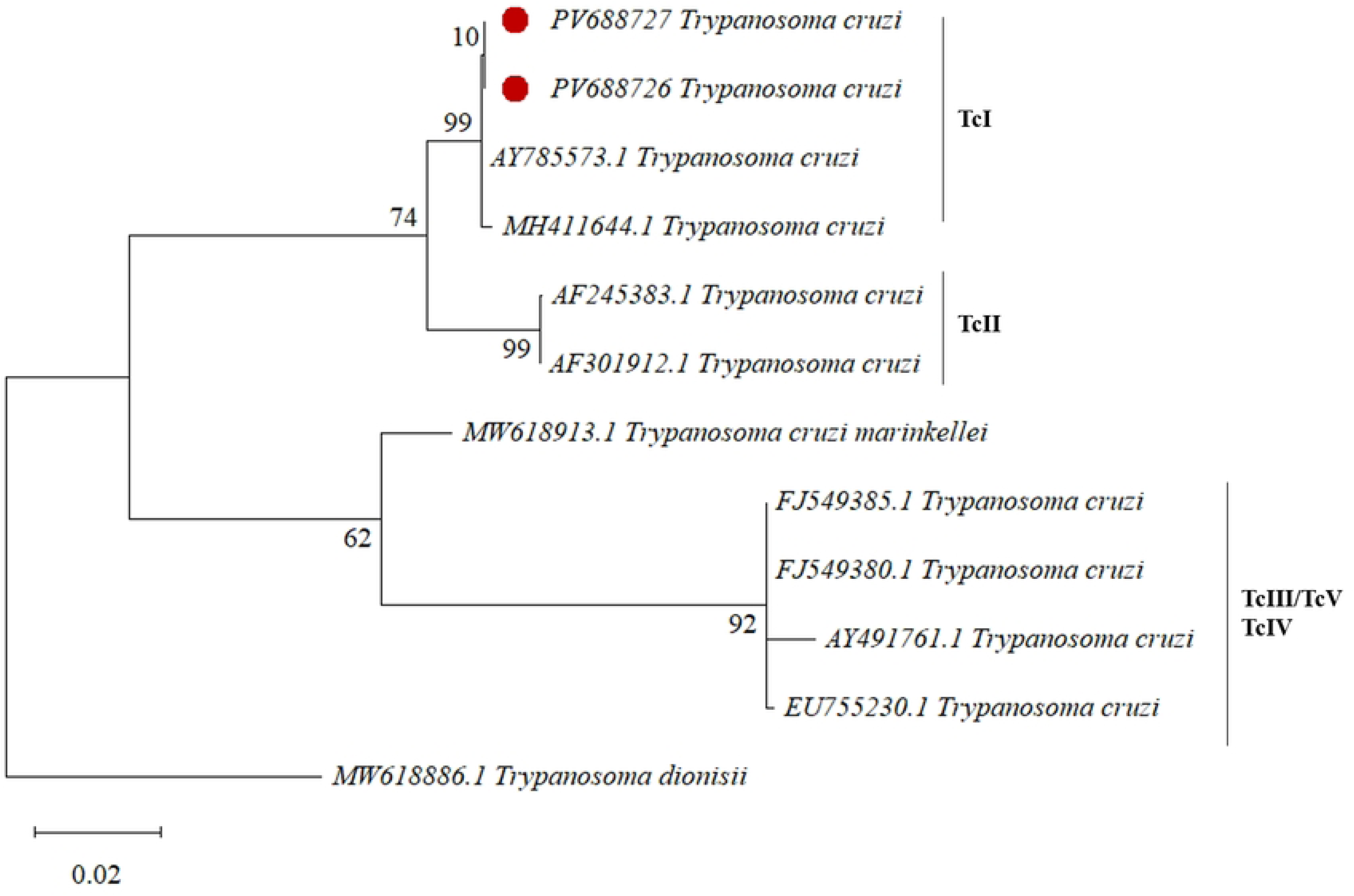
Phylogenetic tree of *Trypanosoma cruzi* based isolates from small mammals captured in Mata da Tapera Municipal Natural Park (MT), Minas Gerais, Brazil. The tree was inferred from 18S SSU rDNA sequences using the Tamura-Nei model with gamma-distributed rate variation among sites (TN+G). Bootstrap values (1,000 replicates) are shown at the nodes. Sequences obtained in this study are marked with red circles and cluster with previously described *T. cruzi* TcI sequences. The scale bar indicates the number of nucleotide substitutions per site.

A total of 102 biological samples, including ear skin, tail skin, spleen, liver, heart, and blood, from 17 individuals were subjected to in vitro culture. Of these, 39 samples (38.2%) were discarded due to contamination, and 60 (58.8%) were negative after three months of monitoring. Trypanosomatids were isolated from three blood samples (2.9%): two from *D. albiventris* and one from *N. squamipes*. Species identification based on 18S rDNA (V7V8 region) nested PCR revealed *T. cruzi* in one *D. albiventris* sample and *T. lainsoni* in both *D. albiventris* and *N. squamipes* (Table 2). Notably, a mixed infection was identified in *N. squamipes* (ID09), with *L. infantum* detected by xenodiagnosis and *T. lainsoni* isolated from blood culture.

**Table 2:** Xenodiagnosis, *in vitro* culture, and 18S rDNA Nested PCR results for small mammals captured in the Mata da Tapera Municipal Natural Park (Minas Gerais, Brazil) between 2023 and 2024.

| ID | Species | Xenodiagnoses<br>(Positive/Total)<br>Accession number |  | Culture<br>(samples) | PCR<br>(samples) |
| --- | --- | --- | --- | --- | --- |
|  |  | *L. longipalpis* | *R. neglectus* |  |  |
| ID01 | *D. albiventris* | *L. infantum*<br>(1/20)<br>PV688719 | Negative<br>(0/10) | Negative | Negative |
| ID02 | *D. albiventris* | Negative<br>(0/20) | *T. cruzi*<br>(5/10) | *T. lainsoni*<br>(blood)<br>PV688723 | *T. lainsoni*<br>(tail skin)<br>PV688724<br>*T. cruzi*<br>(blood and lymph<br>node)<br>PV688726 |
| ID03 | *N. squamipes* | Negative<br>(0/30) | Negative<br>(0/10) | Negative | Negative |
| ID04 | *M. incanus* | *L. braziliensis*<br>(1/10)<br>PV688722 | Negative<br>(0/10) | Negative | Negative |
| ID05 | *N. squamipes* | *L. infantum*<br>(1/20)<br>PV688720 | Negative<br>(0/10) | Negative | Negative |
| ID06 | *N. squamipes* | Negative<br>(0/10) | Negative<br>(0/10) | Negative | Negative |
| ID07* | *M. incanus* | - | - | Negative | Negative |
| ID08** | *D. albiventris* | - | - | - | - |
| ID09 | *N. squamipes* | *L. infantum*<br>(1/46)<br>PV688721 | Negative<br>(0/10) | *T. lainsoni*<br>(blood)<br>PV688725 | Negative |
| ID10* | *M. incanus* | - | - | Negative | Negative |
| ID11 | *M. incanus* | Negative<br>(0/40) | Negative<br>(0/10) | Negative | Negative |
| ID12 | *N. squamipes* | Negative<br>(0/30) | Negative<br>(0/10) | Negative | Negative |
| ID13 | *N. squamipes* | Negative<br>(0/30) | Negative<br>(0/10) | Negative | Negative |
| ID14 | *D. albiventris* | Negative<br>(0/24) | Negative<br>(0/10) | Negative | Negative |
| **ID15** | *N. squamipes* | Negative<br>(0/28) | Negative<br>(0/10) | Negative | Negative |
| **ID16** | *D. albiventris* | Negative<br>(0/30) | *T. cruzi*<br>(2/10) | *T. cruzi*<br>(blood)<br>PV688727 | *T. cruzi*<br>(blood)<br>PV688727 |
| **ID17** | *N. squamipes* | Negative<br>(0/30) | Negative<br>(0/10) | Negative | Negative |
| **ID18** | *N. squamipes* | Negative<br>(0/29) | Negative<br>(0/10) | Negative | Negative |

\* These animals (ID07 and ID10) died before xenodiagnoses; \*\* This specimen of *Didelphis albiventris* was released due to the presence of offspring.

Direct molecular detection using nested PCR targeting the V7V8 region of 18S rDNA confirmed *T. lainsoni* in a tail skin sample and *T. cruzi* in blood and lymph node samples from a single *D. albiventris* individual (ID02), indicating a mixed infection (Fig 5; Table 2). Additionally, *T. cruzi* DTU TcI was detected in the blood of another *D. albiventris* individual (ID16).

**Fig 5:**
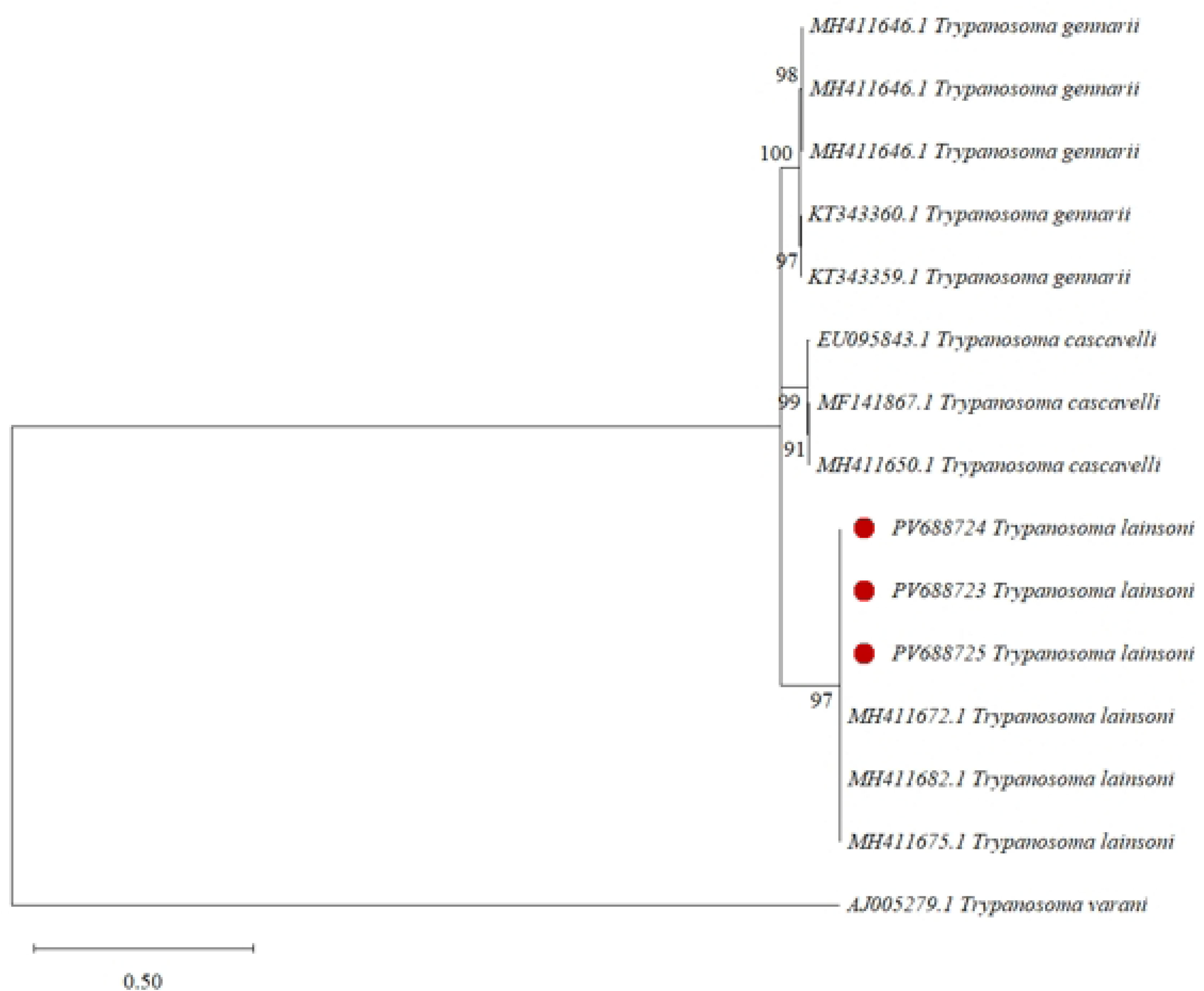
Phylogenetic tree of *Trypanosoma lainsoni* based isolates from small mammals captured in Mata da Tapera Municipal Natural Park (MT), Minas Gerais, Brazil. The tree was inferred from 18S SSU rDNA sequences using the Tamura-Nei model with gamma-distributed rate variation among sites (TN+G). Bootstrap values (1,000 replicates) are shown at the nodes. Sequences obtained in this study are marked with red circles and cluster with previously described *T. lainsoni* sequences from different mammalian hosts. The scale bar indicates the number of nucleotide substitutions per site.

## Discussion

The proximity of Mata da Tapera (MT) to neighboring residential areas facilitates interactions between humans, domestic animals, and wildlife across natural and urbanized interfaces. In fragmented landscapes, common in urban settings, resource scarcity can reduce the viability of larger wildlife populations, leading to biodiversity loss and shifts in host and vector communities [5]. Such environments tend to favor synanthropic species, such as *D. albiventris*, and those with limited home ranges, like *N. squamipes* and *M. incanus* [25,26]. Habitat edges, in turn, intensify contact between humans, vectors, and reservoirs, facilitating the transmission of zoonotic protozoa, especially multi-host pathogens such as *Leishmania* spp. and *Trypanosoma cruzi* [8]. In the present study, one rodent (*N. squamipes*) and two marsupials (*D. albiventris* and *M. incanus*) were naturally infected with trypanosomatids. These findings support the role of small mammals in maintaining parasite cycles in both sylvatic and synanthropic environments [13,27–29]. The detection of zoonotic parasites in mammals from peri-urban areas raises public health concerns due to the potential for spillover to humans and domestic animals.

*Leishmania infantum*, the etiological agent of visceral leishmaniasis (VL) in Brazil, is primarily transmitted by *Lutzomyia longipalpis* [30]. This species was both employed in xenodiagnostic assays and recorded in MT (unpublished data). Although *Lu. longipalpis* is the main vector, *Pintomyia fischeri* and *Migonemyia migonei*, also identified in MT and in neighboring areas such as Jaboticatubas, may act as secondary vectors [15]. Detection of *L. infantum* in three small mammals (11.76%), one *D. albiventris* and two *N. squamipes*, suggests that these species may serve as alternative sylvatic or synanthropic hosts, despite the traditional role of domestic dogs as the principal urban reservoirs of VL. Natural infections in *D. albiventris* have been documented in different regions [5,31,32], reinforcing its relevance at the wildlife–urban interface.

Infection by *L. braziliensis* was detected exclusively through xenodiagnosis in *M. incanus*. Unlike *L. infantum*, typically associated with canids, *L. braziliensis* is more frequently found in hosts from the orders Xenarthra and Rodentia [31]. Its main vector, *Nyssomyia whitmani*, has also been recorded in MT (unpublished data). Previous studies in the Serra do Cipó region identified *Ny. whitmani* carrying *L. braziliensis* and *L. infantum* DNA [15], and in other parts of Minas Gerais, carrying *L. guyanensis* [13], suggesting a broader role in sylvatic cycles of multiple *Leishmania* species.

*Trypanosoma cruzi* DTU TcI was detected in two *D. albiventris* individuals by culture, PCR, and xenodiagnosis. Marsupials of the order Didelphimorphia are recognized for their high infectivity with *T. cruzi*, especially DTU TcI, across multiple Brazilian biomes [28]. Rodents also exhibit high susceptibility and parasitemia levels, contributing as secondary reservoirs [33]. The DTU TcI identified here is widespread in Brazil and strongly associated with transmission by *Rhodnius* spp., consistent with the arboreal habits of both vector and hosts [34]. Although *R. neglectus* was used in xenodiagnoses, field-collected specimens of *Panstrongylus megistus* were also found near palm trees in MT. This species is a well-documented T. cruzi vector in Minas Gerais [16,17], warranting further investigation into its infection status in the study area.

*Trypanosoma lainsoni*, a non-pathogenic species described in 2013, remains poorly characterized [35,36]. Initially placed in the “Megatrypanum” group, it has recently been proposed as part of the LSRM clade, which includes parasites of lizards, snakes, rodents, and marsupials [37]. This study reports the first natural infection of *N. squamipes* by *T. lainsoni*, extending its known host range. No natural vectors have been confirmed for this species. However, related taxa such as *T. (Squamatrypanum) gennarii* have been cultured in different insect species [38]. The failure to detect *T. lainsoni* in *R. neglectus* or *L. longipalpis* suggests vector incompetence, but alternative routes, such as oral transmission through grooming or feeding, cannot be ruled out.

Among all positive mammals (n = 6), only one individual (*D. albiventris*, ID16) tested positive for the same parasite (*T. cruzi*) by all diagnostic methods (culture, PCR, xenodiagnosis). In contrast, *Leishmania* infections were identified exclusively through xenodiagnosis, likely due to low parasitemia. Under such conditions, sand fly infection followed by metacyclogenesis, amplified by euthanizing insects five days post-feeding, may enhance detectability [39]. Furthermore, subsequent PCR targeting 18S rDNA increased sensitivity. While low parasitemia may limit the effectiveness of direct parasitological or molecular methods, it may still suffice for vector infection and transmission in nature. Molecular tools offer high sensitivity and specificity, while culture permits isolation of viable parasites, and xenodiagnosis provides insights into vector competence and host infectiousness [40]. Nonetheless, the scarcity of representative wild mammal samples and the limitations of individual methods remain major constraints in zoonotic disease surveillance. Thus, an integrative diagnostic approach remains essential for reliable detection.

The detection of *L. infantum*, *L. braziliensis*, *T. cruzi* (DTU TcI), and *T. lainsoni* in small mammals from MT underscores the ecological and epidemiological complexity of this peri-urban forest fragment. These findings emphasize the relevance of fragmented landscapes in sustaining parasite transmission cycles, with potential implications for nearby human populations. Integrating parasitological, molecular, and experimental techniques proved effective in identifying natural infections, but future studies should also include vectors and domestic animals, particularly dogs, to clarify their roles in local transmission dynamics. Altogether, our findings reinforce the importance of coupling ecological surveillance with public health actions, especially in areas like MT, where ecotourism, environmental change, and urban encroachment heighten the risk of zoonotic disease emergence.

## Acknowledgments

To the Fiocruz Network of Technological Platforms at Instituto René Rachou - Fiocruz Minas for the DNA sequencing facility, and their assistance and DNA sequencing services.

